# Elucidating Bat SuperImmunity: Insights from the Bent-Wing Bat Genome Assembly and Multi-Omics Analysis

**DOI:** 10.64898/2026.11.11.765934

**Authors:** Carlos Herrera-Ubeda, Michal Zawisza-Alvarez, Maria Catalan-Rabaneda, Jordi Serra-Cobo, Jordi Garcia-Fernandez

**Author notes:** Correspondence to: Carlos Herrera-Ubeda and Jordi Garcia-Fernandez. **Contributions** C.H-U. and J.G-F. contributed with the planning and main ideas of the manuscript; C.H-U. contributed with all the data analysis and main writing of the manuscript; M.Z-A. contributed with the RNA-editing data; M.C-R. contributed with annotation; J.S-C. contributed with dissections, specimen recollection and introduction; All authors contributed to the discussion, corrections and proofreading.

## Abstract

Bats have evolved a range of unique adaptations, including powered flight, echolocation, and exceptional longevity. They have also attracted considerable scientific attention, particularly since the COVID-19 pandemic, due to their association with a broad diversity of viruses and their potential role in zoonotic transmission and emerging infectious diseases. Despite hosting numerous viruses, bats often exhibit limited clinical manifestations of infection, suggesting the evolution of distinctive mechanisms of immune regulation and tolerance. Understanding the molecular basis of these mechanisms has become a major focus of current research, with genome-wide analyses providing important insights into the genetic adaptations underlying bat immunity. Nevertheless, the regulatory complexity of bat genomes, particularly the contribution of non-coding RNAs and post-transcriptional RNA editing, remains comparatively underexplored. Here, we present an integrated multi-omic framework for the common bent-wing bat (*Miniopterus schreibersii*), combining a contiguous chromosome-scale genome assembly with multi-tissue transcriptomic profiling, long non-coding RNA (lncRNA) identification and synteny analysis, endogenous viral elements (EVE) mapping, and same-individual RNA editomic analyses. This study provides a comprehensive resource for investigating genome organization, transcriptional regulation, evolutionary relationships, and post-transcriptional modifications in this species. In particular, it offers an opportunity to explore the potential contribution of non-coding regulatory elements and RNA editing to the molecular mechanisms underlying viral interactions and immune regulation in bats. Collectively, these resources provide a valuable foundation for advancing our understanding of bat genome evolution and the regulatory mechanisms potentially associated with their distinctive responses to viral infections.

## Introduction

The Chiroptera order, more commonly known as bats, is the only group of Mammalia that have achieved powered flight ^1^. This by itself makes bats incredibly interesting creatures, but their uniqueness does not end there. Like cetaceans, they have developed an incredible adaptation such as echolocation ^2^ and they also have a remarkably long lifespan, greater than most mammal species of the same size (for example mice and shrews) ^3^. All these characteristics make the evolution of bats a very successful singular history amongst the mammals, one that has produced an enormous diversity of species (more than 1500 species) adapted to a great spectrum of environments. Bats are the second largest order of mammals, representing 22% of all extant mammalian species ^4,5^ with presence in all the continents except Antarctica ^6^.

But, perhaps, their most remarkable adaptation is their unique relationship with viruses that makes them both the perfect reservoir and vector ^7^. They have been implicated in numerous emerging infectious disease events and have been recognized as important reservoir hosts for viruses that can cross the species barrier and infect humans and other domestic and wild mammals. Their role in viral diseases is well established, particularly for lyssaviruses and coronaviruses ^8–11^. These viruses apparently cause little or no pathology in bats, after years of coevolutionary processes between bats and viruses ^12^. The roles played by bats in the maintenance and transmission of viruses require consideration of the unique characteristics that distinguish bats from all other mammals. They have a high mobility and potential to spread rapidly and widely the viruses. Their gregarious roosting behaviour, particularly during the summer season, where they closely cluster together, can significantly increase the likelihood of viral transmission within the same species and between different bat species ^13^. The aforementioned extreme longevity of bats, together with the possibility that they might develop persistent infections with certain viruses, may help to maintain the viruses and transmit them to other vertebrates ^13^. These characteristics make the study of their genome a crucial element. This is not only true for understanding their evolution, but for trying to infer how they achieved their unique viral-host relationship.

Recently, research on bats has seen a surge of exciting new data, with new genomes that have been already sequenced and analysed ^14^, and new transcriptomics with an extensive amount of work done in comparative genomics. But most of current bibliography regarding these topics though, is focused on coding genes, with only a handful of works versing on non-coding RNAs for example ^15^. In this regard, initiatives like the bat1k project ^16^ are helping to elucidate more and more how the genomic features of these species are organised. One of the most gregarious bat species in Southern Europe, with one of the greater lifespans observed in our studies (19 years), is Schreiber’s Bent-winged Bat (*Miniopterus schreibersii*) ^17–19^. This bat species has been recently described as capable of harbouring and spreading coronaviruses ^20^ but has yet to be analysed thoroughly at the genomic level. *M. schreibersii* represents a specialized evolutionary lineage (family Miniopteridae) characterized by long, narrow wings adapted for fast hawking flight in open spaces, extensive seasonal migrations, cave-dwelling nursery and winter colonies numbering tens of thousands of individuals. These ecological traits place *M. schreibersii* in a position of elevated viral exposure.

Here we present a newly assembled Schreiber’s Bent-winged Bat genome of an individual from Catalonia (Spain), with chromosome level assembly and a comprehensive analysis of chromosome architecture, lncRNAs, EVEs, and RNA-editing, focused on the evolutionary differences with reference mammals.

## Results

### Genome Data and Chromosome-Scale Architecture

Previous assemblies of *M. schreibersii* (GCA_004026525.1, GCF_964146895.1) suggested a genome size of around 1784 Mb distributed in 22 chromosomes ^21^. Our assembly, however, has a slightly lower size with around 1767 Mb distributed in 23 different chromosomes after HiC chromosome assembly, with the Y chromosome fragmented in four heterochromatic scaffolds totalling 4.05Mb (Table 1, Figure 1). Interestingly there are 3 micro-chromosomes harbouring contacts between themselves as well as with the centromeric regions of other chromosomes (Figure 1). The N50 nanopore reads after removing outliers had a size of 34.33 kb with 116x nuclear coverage. The mitochondrial genome of *M. schreibersii* was assembled into a single circular contig of 16,640 bp presenting two replication origins and 37 genes (13 protein-coding genes, 2 rRNAs, and 22 tRNAs). The three smallest autosomes, SUPER_20 (27.86 Mb), SUPER_21 (17.96 Mb), and SUPER_22 (11.97 Mb), exhibit distinctive structural and compositional characteristics (Supplementary Table 1). Compared to macrochromosomes (SUPER_1 to SUPER_9), which have a mean GC content of 41.3% and a mean gene density of 42.4 transcripts/Mb, these three chromosomes possess elevated GC content (46.53% to 49.67%), the lowest repeat content (22.2%–23.5%), and the highest transcript densities across the assembly (61.5 to 87.8 transcripts/Mb). They encode 4523 annotated transcripts, including essential genes (CHEK1, IL16), implying that they are euchromatic A-chromosomes rather than dispensable B-chromosomes.

**Fig. 1:**
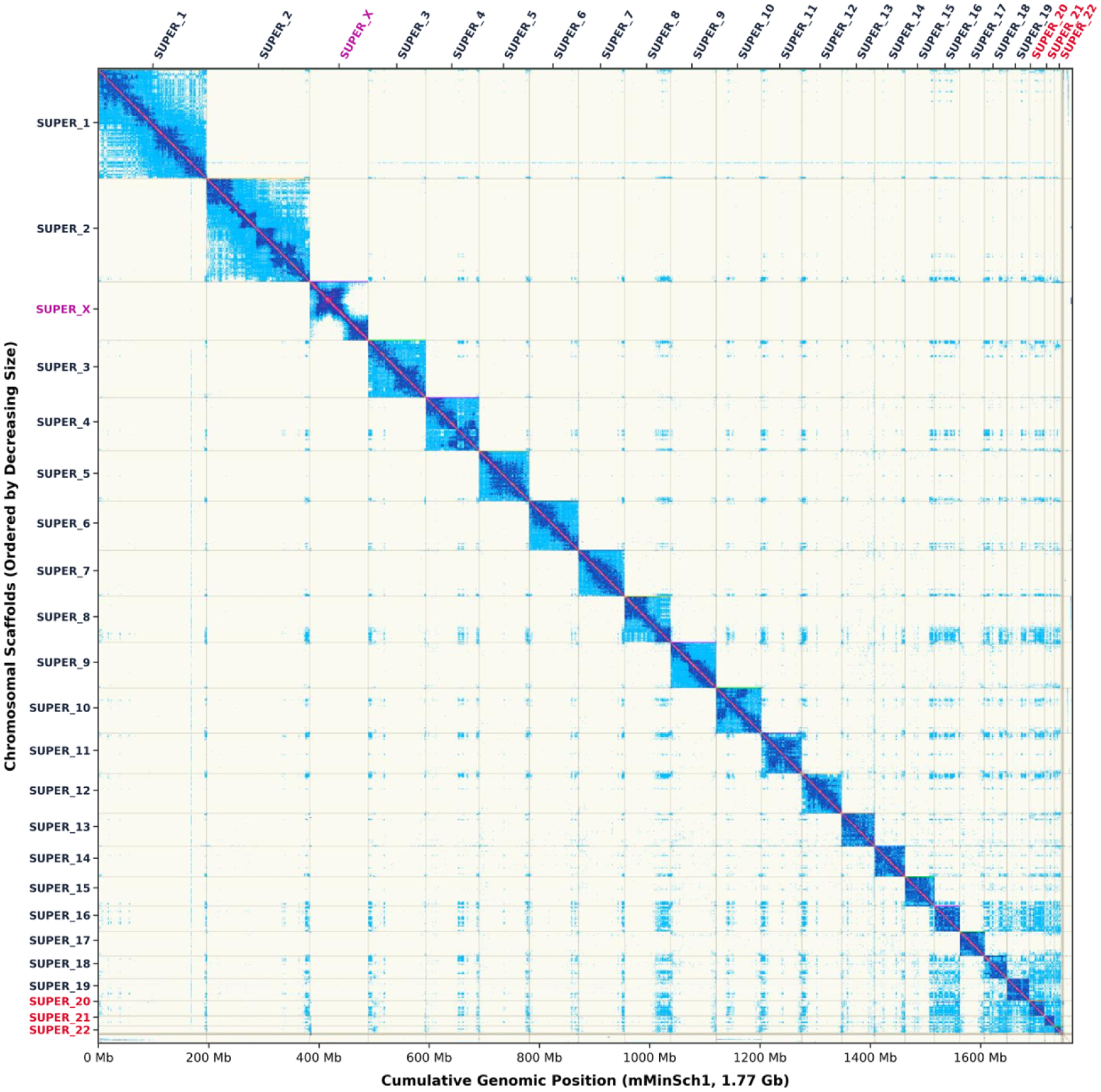
Genome-wide Hi-C chromatin contact map. Genome-wide Hi-C chromatin contact map of the chromosome-level *M. schreibersii* assembly ordered by decreasing length with the three smallest autosomes marked in red, the X chromosome in purple, and th fragmented SUPER_Y and unplaced scaffolds in the bottom-right margin.

**Table 1.** Genome assembly traits: Expected vs. Observed metrics in *M. schreibersii*.

| **Genome Traits** | **Expected** | **Observed** |
|---|---|---|
| **Haploid size (bp)** | 1,784,288,419 | 1,767,139,219 |
| **Haploid Number** | 22 | 23 |
| **Ploidy** | 2 | 2 |
| **Sample Sex** | XY | XY |

Regarding the BUSCO analysis, we found 98.2% of completeness, (96.7% single copy and 1.5% of duplicates) with 0.8% of fragmented hits and a 1% of missing hits. The complete summary of all the genomic features can be found in Supplementary Table 1. The final curated assembly achieved the Earth BioGenome Project (EBP) quality standard of 7.7.Q53 (consensus quality value QV = 53.29, k -mer completeness = 98.1%).

Using RNA extraction from different tissues (Wing epithelium, heart, liver, intestines, spleen, brain, lungs, testis and muscle) we performed RNAseq and assembled a transcriptome with 86663 transcripts. Of those, 67948 consisted of protein coding CDS with a BUSCO completeness of 92.1%. Finally, when running orthofinder using 20 vertebrate proteomes (see methods), a total of 17405 orthogroups were found.

### Gene Family Expansions and Contractions

Analysing the expansions and contractions of gene families in *M. schreibersii* against a set of 20 vertebrate proteomes with 9 chiropterans, 6 non-chiropteran laurasiatherians, two euarchontoglirans, one afrotherian, one marsupial metatherian and one non-mammalian amniote (acting as outgroup; see methods) raw CAFE5 modelling found 151 families expanded and 565 contracted (p-value < 0.05). After filtering the expansions to counter annotation artifacts, we ended with 78 bona fide expansions within the *Miniopterus* genome (Supplementary Table 2).

Enrichment analysis of the 78 curated expanded families revealed overrepresentation of the Hedgehog signalling pathway (the only KEGG term significant after FDR correction) along with canonical Wnt regulation, intracellular iron sequestration and protection against ferroptosis, as well as positive regulation of DNA repair and SWI/SNF chromatin remodelling. Conversely, contracted families were heavily dominated by olfactory receptors, reflecting the evolutionary sensory trade-off favouring echolocation over olfaction, alongside sensory and inflammasome sensor modulations (Figure 2). Some of the most interesting expansions and contractions as well as the proposed biological reasoning can be found summarized in Table 2.

**Fig. 2:**
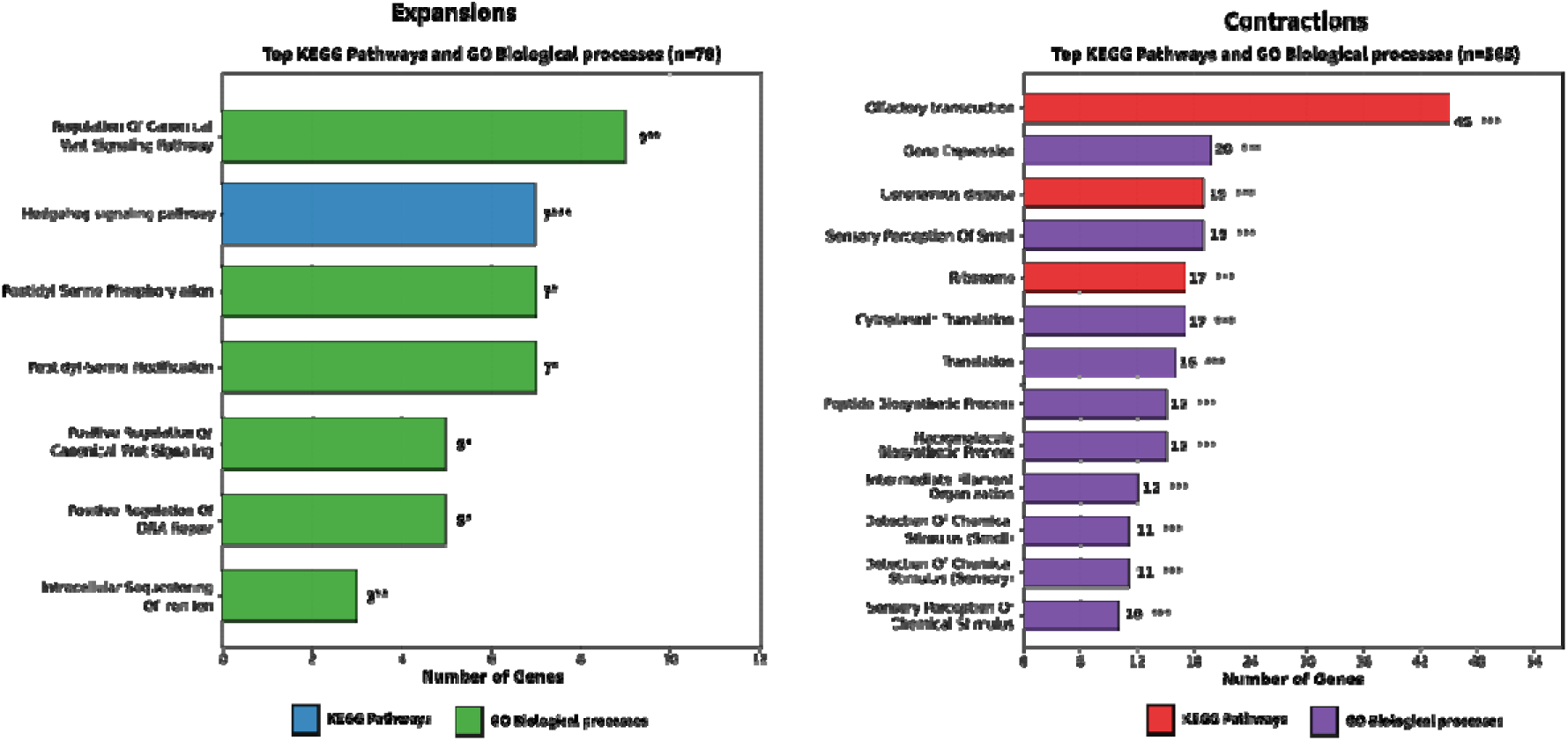
Functional enrichment of curated gene family dynamics in *M. schreibersii*. Horizontal bars represent the number of genes identified per category across KEGG pathways and Gene Ontology Biological Processes for curated expansions (n = 78) and contractions (n = 565). Significance (Enrichr Fisher exact test, Benjamini-Hochberg adjusted p, FDR) is indicated at the tip of each bar (* FDR < 0.05, ** FDR < 0.005, *** FDR < 0.0005). Gene counts refer to properly annotated genes (unnamed Hypcod/LOC loci cannot be mapped to the Enrichr libraries).

**Table 2.** CAFE5: Curated Key Gene Family Expansions and Contractions in *M. schreibersii*.

| Orthogroup ID | Direction | Net Change | Branch p-value | Representative Genes (*M. schreibersii*) | Functional Category / Biological Significance |
|---|---|---|---|---|---|
| **OG0000264** | Expansion | +14 | < 0.0001 | *ACTL6B*, *Hypcod01927*, *Hypcod02316* | **Neural Plasticity & Epigenetics**: Subunit of neural SWI/SNF (BAF) chromatin remodeling complex; synaptic plasticity and long-term memory |
| **OG0001057** | Expansion | +6 | < 0.0001 | *CSNK1A1*, *CSNK1D*, *CSNK1E*, *CSNK1G1-3* | **Cellular Signaling & Circadian Rhythm**: Casein kinase 1 family; regulation of Wnt, Hedgehog, and circadian pacemaking |
| **OG0000080** | Expansion | +5 | < 0.0001 | *FTH1*, *FTL2*, *FTL3*, *FTL4*, *FTMT* | **Flight Metabolism & Stress Tolerance**: Iron sequestration and antioxidant protection against ROS from intense flight metabolism |
| **OG0000323** | Expansion | +5 | < 0.0001 | *RALGDSL*, *RALGDSL2*, *LOC03985* | **Autophagy & Tumor Suppression**: Ral-GEF signaling; cellular stress response, genomic maintenance, and longevity |
| **OG0000139** | Expansion | +4 | < 0.0001 | *GIMAP1*, *GIMAP2*, *GIMAP5B-D* | **Lymphocyte Homeostasis**: GTPases of immunity-associated proteins; regulation of T- and B-cell survival and apoptosis |
| **OG0001229** | Expansion | +4 | < 0.0001 | *CCDC88C* (Daple), *Hypcod00720* | **Epithelial Integrity & Polarity**: Non-canonical Wnt transducer; cell junction integrity and mechanical elasticity in wing patagium |
| **OG0000101** | Expansion | +3 | 0.0001 | *TRIM60*, *TRIM61*, *TRIM75* | **Antiviral Defense**: Tripartite motif (TRIM) E3 ligases; restriction of retroviruses and enveloped RNA viruses |
| **OG0000094** | Expansion | +2 | 0.0035 | *HSPA1L*, *HSPA8* | **Flight Heat Shock Tolerance**: Hsp70 chaperones buffering flight-induced hyperthermia and proteotoxic stress |
| **OG0001359** | Expansion | +2 | 0.0001 | *TPM3B*, *TPM3D*, *TPM4* | **Muscle Biomechanics**: Tropomyosins; sarcomeric actin regulation supporting high wingbeat frequency |
| **OG0000005** | Contraction | -9 | < 0.0001 | *OR4A47*, *OR4C11*, *OR4C15*, *OR4C3* | **Sensory Trade-off**: Olfactory receptor family 4 contraction (echolocation reliance) |
| **OG0000006** | Contraction | -8 | < 0.0001 | *OR51B5*, *OR51B6*, *OR51G1-2* | **Sensory Trade-off**: Olfactory receptor family 51 contraction |
| **OG0000017** | Contraction | -6 | < 0.0001 | *OR13C8*, *OR13F1*, *OR13J1*, *OR2K2* | **Sensory Trade-off**: Olfactory receptor families 13 and 2 contraction |
| **OG0000036** | Contraction | -6 | < 0.0001 | *NLRP1*, *NLRP5*, *NLRP14* | **Anti-inflammatory Adaptation**: Contraction of NLRP inflammasome sensors; attenuation of cytokine storm and hyperinflammation |
| **OG0000016** | Contraction | -5 | < 0.0001 | *OR10J3*, *OR10W1*, *OR10Z1* | **Sensory Trade-off**: Olfactory receptor family 10 contraction |
| **OG0000009** | Contraction | -4 | < 0.0001 | *OR52B2*, *OR52B4*, *OR52D1*, *OR52E2* | **Sensory Trade-off**: Olfactory receptor family 52 contraction |
| **OG0000024** | Contraction | -3 | < 0.0001 | *IFNB1*, *IFNK* | **Interferon Modulation**: Type I interferon locus contraction (restructured baseline antiviral state) |
| **OG0000011** | Contraction | -3 | 0.0005 | *LILRA5*, *LILRA6*, *NCR1* | **Innate NK Regulation**: Leukocyte immunoglobulin-like receptor & Natural cytotoxicity receptor |
| **OG0000015** | Contraction | -8 | < 0.0001 | *MR1* | **Antigen Presentation**: MHC class I-related protein contraction |
| **OG0000008** | Contraction | -10 | < 0.0001 | *MAGEA10*, *MAGEA11*, *MAGEB16*, *MAGEB3* | **Germline / Differentiation**: Melanoma antigen gene family |

### Endogenous Viral Elements (EVEs) & Tissue Expression Profiles

Using detectEVE against the chromosome-level assembly (see methods) to find the Endogenous Viral Elements within the genome of *M. schreibersii*, we found a total of 433 high confidence hits to the default detectEVE databases (Supplementary Table 3). We included in the analysis both dsDNA and retroviruses, with further filtering to avoid false positives. From the 433 EVEs, most of them were classified as Artverviricota or Peploviricota with a small subset of 21 non-retroviral EVEs (nrEVEs).

This subset of nrEVEs was subjected to further analysis by strict domain curation (Pfam-A) and comparative synteny analysis with *M. myotis* in order to avoid false positives. 14 of them were found to be artifacts due to gene acquisition events, such as EVE169 which directly overlaps the third exon and second intron of host interleukin-10 (IL10) on SUPER_19. As shown by Irwin et al. 2021, Nucleocytoviricota are known to regularly encode “cellular like” genes to manipulate host defences among others ^23^. After removing these 14 artifacts we validated seven bona fide nrEVEs possessing canonical viral protein domains and normal sequence complexity (36.7%–52.4% GC): four Negarnaviricota elements (two Mononegavirales L-polymerase integrations and two Bornaviridae P40 nucleoprotein integrations), two X-linked Cossaviricota (Parvoviridae NS1/Rep) elements, and one Preplasmiviricota element (Table 3).

**Table 3.** Comprehensive Expression Profile of the 7 curated nrEVEs in *M. schreibersii*.

| Genomic Coordinate | EVE ID | Viral Phylum (Family / Protein) | Pfam Domain | Feature Overlap | Host Gene ID | Mean TPM (All 24 Samples) | Primary Tissue of Expression (Max TPM) | Expressed Samples Count (>0.1 TPM) |
|---|---|---|---|---|---|---|---|---|
| SUPER_19:33,112,095–33,113,455 | EVE029 | Negarnaviricota (Mononegavirales L) | Mononeg_RNA_pol (PF00946) | intron | LRIG2 | 14.72 | heart_1 (51.29 TPM) | 24 / 24 |
| SUPER_3:27,623,119–27,623,522 | EVE204 | Negarnaviricota (Bornaviridae N) | BDV_P40 (PF06407) | exon (3' UTR) | TBL2 | 6.00 | testis_3 (17.80 TPM) | 23 / 24 |
| SUPER_1:54,440,809–54,441,111 | EVE236 | Preplasmiviricota | Viral ORF (101 aa) | intron | BLOC1S6 | 0.34 | liver_2.2 (1.53 TPM) | 10 / 24 |
| SUPER_10:62,909,917–62,913,457 | EVE003 | Negarnaviricota (Mononegavirales L) | Mononeg_RNA_pol / mRNAcap | Intergenic | — | 0.01 | intestine_3 (0.32 TPM) | 1 / 24 |
| SUPER_14:12,134,448–12,134,993 | EVE098 | Negarnaviricota (Bornaviridae N) | BDV_P40 (PF06407) | Intergenic | — | 0.00 | Undetected (0.00 TPM) | 0 / 24 |
| SUPER_X:74,393,355–74,393,785 | EVE078 | Cossaviricota (Parvoviridae NS1) | Parvo_NS1 (PF01057) | Intergenic | — | 0.00 | Undetected (0.00 TPM) | 0 / 24 |
| SUPER_X:74,393,638–74,394,142 | EVE079 | Cossaviricota (Parvoviridae NS1) | Parvo_NS1 (PF01057) | Intergenic | — | 0.00 | Undetected (0.00 TPM) | 0 / 24 |

Quantifying their transcriptional activity across 24 RNA-seq tissue libraries revealed a sharp functional dichotomy between intergenic and host-gene-integrated nrEVEs (Figure 3, Table 3, Supplementary Table 4). All four intergenic nrEVEs are transcriptionally silenced across bat tissues (0.00–0.01 mean TPM). Conversely, all three nrEVEs integrated within host transcriptional units are actively expressed:

1. **EVE204 (Bornaviridae N/P40 in the 3’ UTR exon of *TBL2*):** Located on SUPER_3 with a length of 404 bp, this element encodes a Bornavirus P40 nucleoprotein domain integrated directly within Exon 7 (the 3’ UTR exon) of host *TBL2*. Comparative alignment with *M. myotis* confirmed high conservation of the flanking *TBL2* locus (86.3%–86.8% nucleotide identity) framing a clean empty pre-integration site in *M. myotis*, exposing a lineage-specific Bornaviridae insertion into the *Miniopterus TBL2* 3’ UTR. De novo transcript assembly (STRG.5278.1) confirmed direct exonic read coverage co-transcribing EVE204 within mature *TBL2* mRNA across 23 of the 24 tissue libraries (mean = 6.00 TPM), peaking in testis, lung, and intestine (testis_3: 17.80 TPM; lung_3: 14.90 TPM; intestine_3: 11.49 TPM).
2. **EVE029 (Mononegavirales L-polymerase in the intron of *LRIG2*):** Spanning 1,361 bp on SUPER_19, this element encodes a Mononegavirales RNA-dependent RNA polymerase domain integrated within the 5’ intron of *LRIG2*. Unlike the *Miniopterus*-specific EVE204, EVE029 is shared at the orthologous intronic position in *M. myotis* (81.1% nucleotide identity), dating its endogenization to the common ancestor of Vespertilionoidea (> 45 MYA). The host *LRIG2* transcriptional unit is ubiquitously expressed across all 24 tissue samples (mean = 14.72 TPM), peaking in cardiac and skeletal muscle (heart_1: 51.29 TPM; heart_3: 43.30 TPM; muscle_3: 27.18 TPM).
3. **EVE236 (Preplasmiviricota in the *BLOC1S6–SQOR* locus):** With a size of 303 bp on SUPER_1, this EVE contains a 101-aa viral-like ORF embedded within a long transcriptional unit (STRG.721) spanning the *BLOC1S6–SQOR* interval. Comparative alignment against the orthologous block in *M. myotis* (scaffold_m19_p_1:163,453,032–163,453,334) revealed that while the surrounding 20 kb of intronic sequence diverges at neutral rates (81.4%–83.7% identity), the 303-bp EVE236 sequence exhibits an increased 96.04% nucleotide identity (*E* = 4.95 x 10^−142^), indicating purifying selection across > 45 million years. Transcriptional profiling showed selective expression across 10 / 24 samples (mean = 0.34 TPM), predominantly in hepatic libraries alongside spleen and intestine (liver_2.2: 1.53 TPM; liver_1.2: 1.25 TPM; liver_2.1: 1.18 TPM; spleen_3: 0.90 TPM; intestine_2.2: 0.61 TPM).

**Fig. 3:**
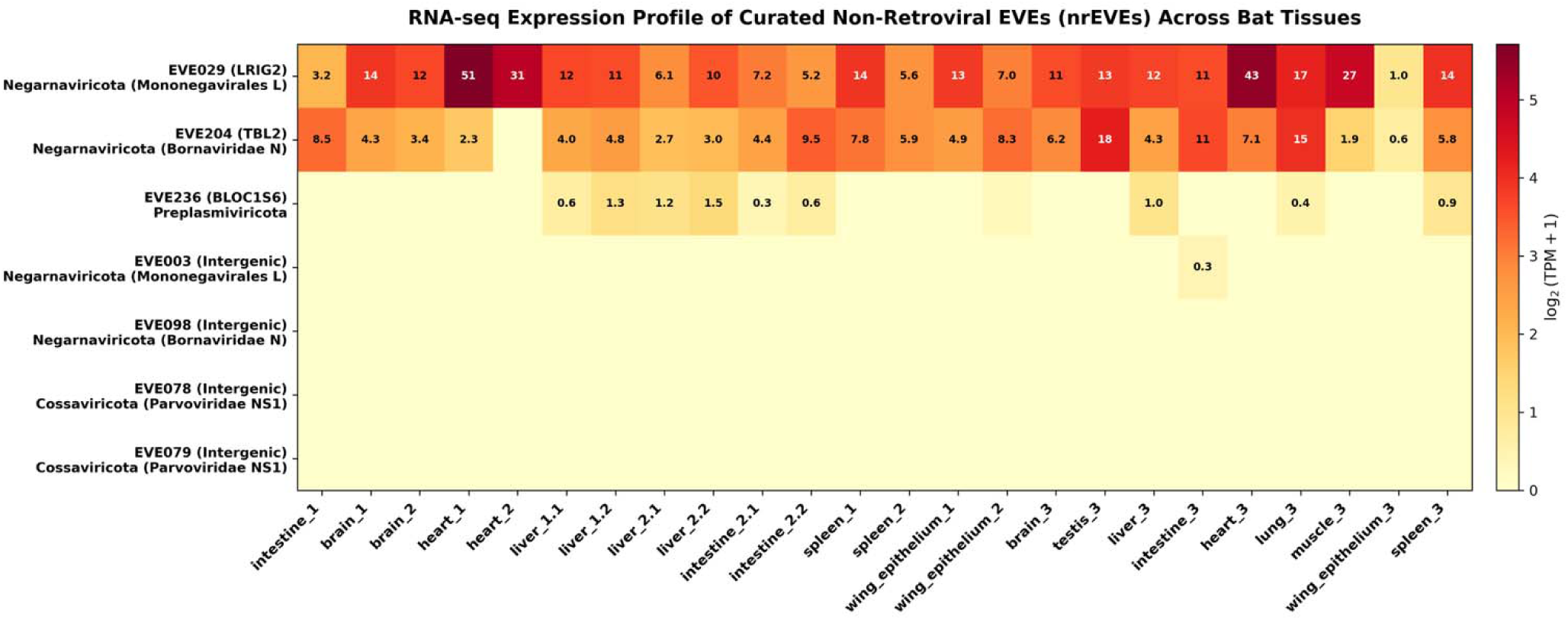
Expression heatmap of the 7 curated nrEVEs identified in available tissues from three bats.

### lncRNA Mining and Positional Microsynteny in Immune Loci

When analysing the non-coding fraction of the *M. schreibersii* genome, we decided to focus specifically on the long noncoding RNAs (lncRNAs). After applying several stringent computational filters to discard all the transcripts with any trace of coding identity using different tools (CPAT, BLAST, HMMER; see methods) we found a total of 7693 putative lncRNAs (Supplementary Table 5). If we classify them regarding their relative position to other coding genes, we observe that, as expected, most of these lncRNAs are classified as Intergenic, with 4433 falling into this category with a great overrepresentation of antisense lncRNAs as well (Figure 4).

**Fig. 4:**
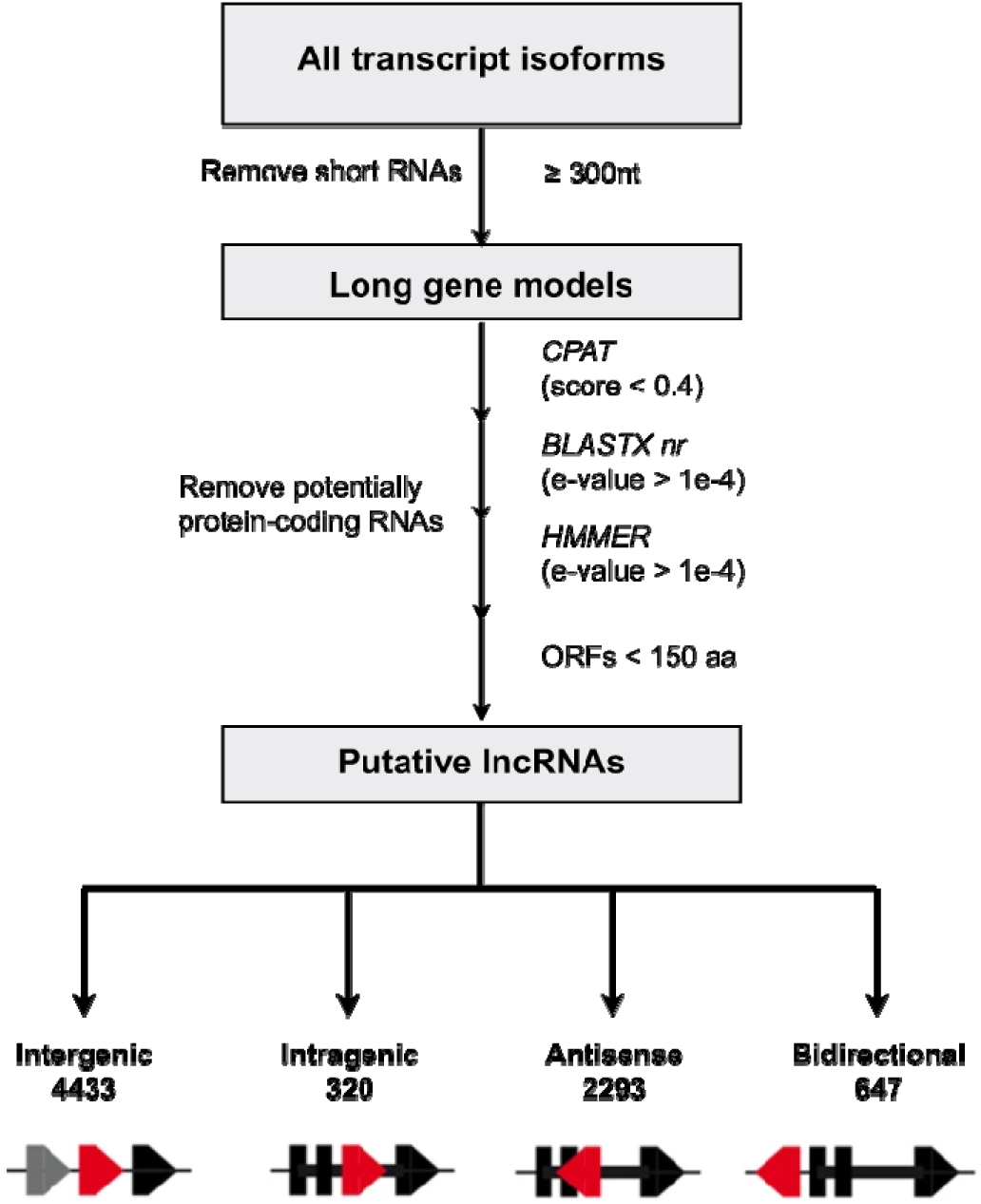
lncRNA pipeline identification and classification.

Using lincOfinder (see methods) between *M. schreibersii* and *H. sapiens*, we determined that 4514 of the 5080 lincRNAs (intergenic and bidirectional) reside in conserved microsyntenic blocks. Additionally, 4106 of these blocks harbour a putative orthologous lincRNA in H. sapiens. Cross-referencing these results against curated immune databases revealed that 360 (8.77%) of the 4106 lincRNAs are flanked by an immune related synteny block. This represents a statistically significant 1.32-fold enrichment on immunity related blocks over genomic background (p-value = 9.59 x 10^-8).

These not only include some well-known conserved lincRNAs such as *HOTAIRM1* ^24^ (Hypnc11349 within the HOXA cluster on SUPER_9) but also conserved lincRNAs flanking the interleukin-4/21 receptor locus, the interferon receptor complex, Toll-like receptor 9, interleukin-1 receptor antagonist, and the B-cell receptor-like locus among others (Table 4).

**Table 4.** Representative Microsyntenically Conserved lincRNAs in Immune Loci.

| lncRNA ID | Bat Coordinate | Syntenic Block (Human Homolog) | Syntenic Immune Genes | Flanking Gene Neighborhood (Syntenic Block) | Immune Subsystem / Pathway |
|---|---|---|---|---|---|
| Hypnc11349 | SUPER_9:72,886,512–72,890,519 | Chr7:26,346,443–27,110,640 (HOTAIRM1) | HOXA1, HOXA2, HOXA9 | HOXA1, HOXA2, HOXA3, HOXA4 | **Myeloid Differentiation**: Intergenic within HOXA cluster; deep mammalian conservation |
| Hypcod01757 | SUPER_3:4,478,540–4,481,350 | Chr16:60,042–27,778,629 (LOC124903617) | IL21R, IL4R | GTF3C1, IL21R, IL4R, KIAA0556, RHBDF1, MPG | **Cytokine Signaling**: IL-4 and IL-21 dual-receptor locus modulating humoral immunity |
| Hypnc09139 | SUPER_5:87,406,912–87,410,535 | Chr21:33,241,923–44,873,616 (IL10RB-DT) | IFNAR1, IFNAR2, IL10RB | IFNAR1, IFNAR2, IL10RB, GART, ITSNA | **Antiviral Cytokine Receptor**: Master type I interferon and IL-10 receptor complex |
| Hypnc03897 | SUPER_15:50,905,679–50,908,062 | Chr3:4,351,527–52,225,529 (ITPR1-DT) | TLR9 | TLR9, ALAS1, SUMF1 | **Innate Sensing**: Intracellular DNA sensing receptor triggering type I interferon |
| Hypnc03134 | SUPER_13:49,659,755–49,660,236 | Chr2:88,067,865–113,132,871 (LOC101928347) | IL1F10, IL1RN | IL1F10, IL1RN, FABP1, SMYD1 | **Anti-inflammatory**: Antagonist of interleukin-1 pro-inflammatory signaling |
| Hypnc05220 | SUPER_19:4,988,729–4,989,529 | Chr1:145,960,269–157,820,037 (LINC01719) | FCRL1 | Hypcod01286, FCRL1, HJV, TXNIP, ANKRD34A, POLR3GL | **B-cell Homeostasis**: Fc receptor-like 1 regulator of B-cell receptor signaling |
| Hypnc05977 | SUPER_2:81,659,716–81,660,092 | Chr13:23,330,136–46,892,513 (SACS-AS1) | TNFRSF19 | TNFRSF19, MIPEP, SACS, HTR2A, ESD, LRCH1 | **TNF Superfamily**: Tumor necrosis factor receptor superfamily signaling |
| Hypnc08456 | SUPER_4:75,347,787–75,348,107 | Chr1:78,688–248,824,935 (LOC124903816) | TRIM58 | ZNF692, TRIM58, RNF187 | **Antiviral Restriction**: Tripartite motif E3 ubiquitin ligase regulation |
| Hypnc10828 | SUPER_8:78,926,178–78,928,317 | Chr1:12,104,444–13,615,911 (LINC02766 / LINC01784) | TNFRSF1B, TNFRSF8 | PDPN, LRRC38, VPS13D, TNFRSF1B, TNFRSF8 | **Immune Checkpoints & Costimulation**: TNF receptor 2 (TNFRSF1B) and CD30 (TNFRSF8) regulatory locus |
| Hypnc05222 | SUPER_19:5,447,380–5,447,702 | Chr1:54,191–159,837,311 (LOC124903816) | CD1D, FCRL2, FCER1A, SLAMF8 | Hypcod01288, CD1D, FCRL2, FCER1A, CADM3, SLAMF8 | **Antigen Presentation & Lymphocyte Activation**: NKT lipid antigen presentation (CD1D) and Fc/SLAM receptors |
| Hypnc05095 | SUPER_18:34,980,356–34,980,670 | Chr16:50,697,244–53,500,259 (CYLD-AS1 / LINC02168) | NOD2, CYLD | SALL1, TOX3, CYLD, NOD2, AKTIP | **Innate Sensing & NF-κB Brake**: Cytosolic pattern sensor NOD2 and NF-κB/IFN deubiquitinase CYLD |
| Hypcod01761 | SUPER_3:5,681,566–5,684,620 | Chr16:20,627,086–28,595,930 (LOC105371121) | IL27 | IL27, APOBR, Hypcod01767, ACSM1B | **Immunoregulatory Cytokine**: Interleukin-27 locus restraining excessive Th1/Th17 inflammation |

### A-to-I RNA-Editing Identification, Functional Recoding, and Deep Learning Cross-Testing

Using RNAseq data from the same individual as the genomic data, we were able to identify SNPs supporting A-to-I RNA editing. This yielded 234736 high-confidence unique putative RNA-editing events (Supplementary Table 6)

Annotation against the curated *M. schreibersii* annotation showed that the vast majority of editing sites reside in non-coding regions: 65.28% (153,227 sites) are intronic and 16.19% (38,012 sites) are intergenic, reflecting the editing of inverted repeat elements (SINEs/LINEs) in primary transcripts to suppress aberrant dsRNA-sensing. Within mature transcripts, 10.79% (25,319 sites across 4,307 genes) mapped to 3’ UTRs, 3.64% (8,535 sites) to 5’ UTRs, and 1.99% (4,664 sites) to non-coding transcript exons (Figure 5A; Table 5; Supplementary Table 6).

**Fig. 5:**
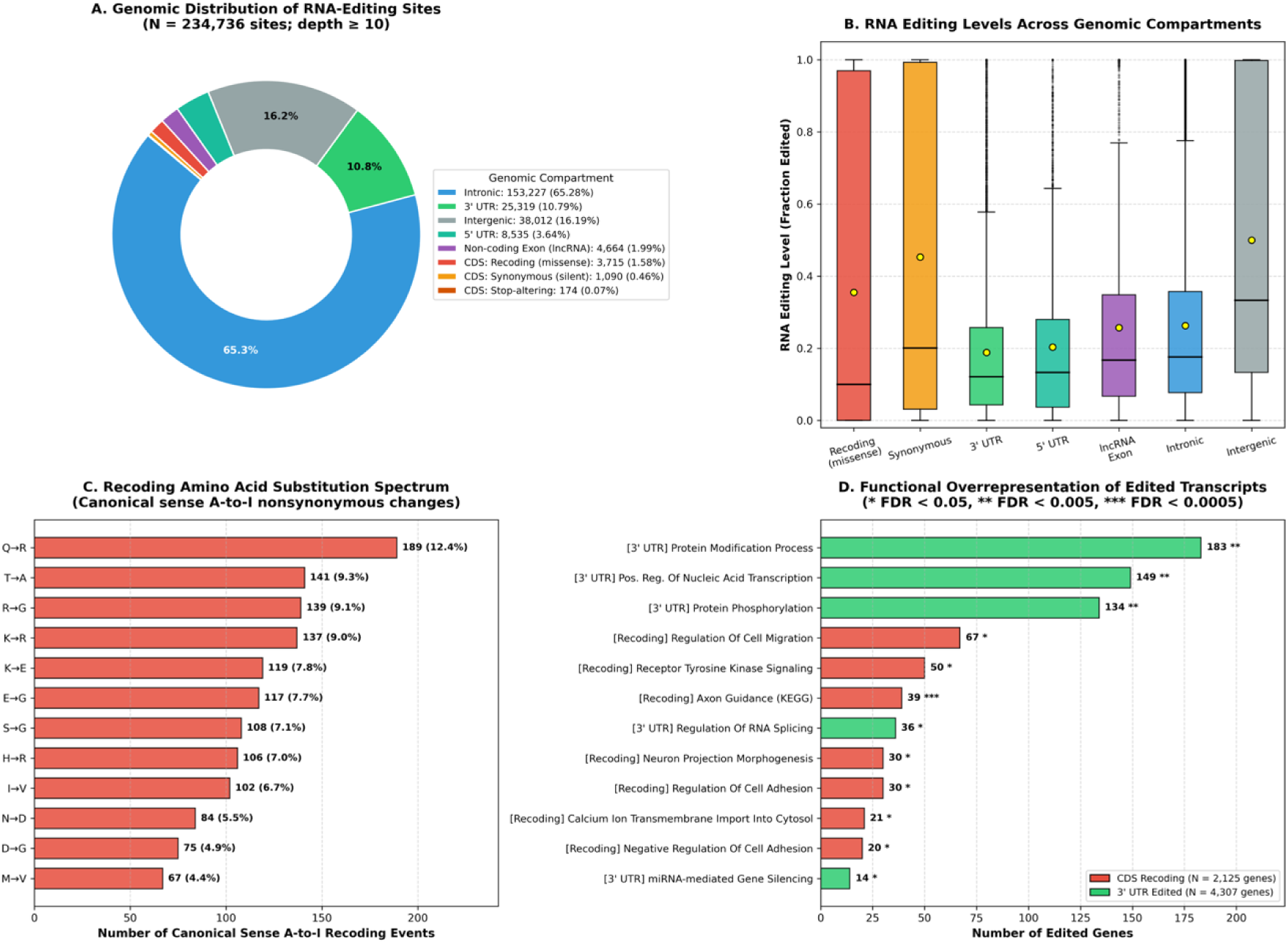
Comprehensive landscape of the *M. schreibersii* RNA editome. **A** Genomic distribution of 234,736 high-confidence RNA-editing sites (depth ≥ 10). Proportions and absolute counts are shown for introns, intergenic space, 3’ UTRs, 5’ UTRs, non-coding transcript exons, CDS recoding events, CDS synonymous sites, and stop-altering positions. **B** Distribution of RNA editing levels (fraction edited) across genomic compartments. Boxes indicate interquartile ranges (IQR), bold horizontal lines represent medians, yellow diamonds denote means, and whiskers delineate 1.5 x IQR. **C** Amino acid substitution spectrum of nonsynonymous CDS editing events, highlighting the dominance of Q->R, T->A, R->G, and K->R transitions. **D** Functional overrepresentation analysis of transcripts undergoing nonsynonymous recoding (red) and 3’ UTR editing (green) across GO Biological Process and KEGG pathways. Bars represent the number of edited genes; significance is indicated at the tip of each bar (FDR < 0.05, ** FDR < 0.005, *** FDR < 0.0005).

**Table 5.** Global RNA Editome Summary Statistics.

| **Genomic Compartment** | **Site Count** | **Percentage of Total Editome (%)** | **Mean Editing Level (%)** |
|---|---|---|---|
| Intronic | 153,227 | 65.28% | 26.3% |
| Intergenic | 38,012 | 16.19% | 50.0% |
| 3' UTR | 25,319 | 10.79% | 18.9% |
| 5' UTR | 8,535 | 3.64% | 20.3% |
| Non-coding Exon (lncRNA) | 4,664 | 1.99% | 25.8% |
| CDS: Recoding (missense) | 3,715 | 1.58% | 35.5% |
| CDS: Synonymous (silent) | 1,090 | 0.46% | 45.3% |
| CDS: Stop-altering | 174 | 0.07% | 66.5% |
| TOTAL EDITOME (depth ≥ 10) | 234,736 | 100.00% | 29.3% |

A total of 4,979 editing events mapped within open reading frames (CDS). In silico translation revealed that 3,715 sites (74.6% of CDS events; 1.58% of total editome) cause nonsynonymous amino acid substitutions (recoding events) distributed across 2,125 unique genes (Supplementary Table 7), while 1,090 sites (21.9%) are synonymous, and 174 sites (3.5%) alter stop codons. Across the editome, editing levels displayed a broad distribution (overall mean = 29.3%, median = 18.2%; Figure 5B), with recoding sites exhibiting an average editing frequency of 35.5% (median = 10.0%, reaching 100% in key physiological targets) and 3’ UTR sites showing an average of 18.9% (median = 12.1%). The recoding amino acid spectrum was dominated by Q→R (12.4%), T→A (9.3%), R→G (9.1%), K→R (9.0%), and K→E (7.8%) substitutions (Figure 5C).

The canonical mammalian recoding site, the Q/R site (Q607R) within the channel pore of the AMPA glutamate receptor subunit GRIA2 ^25–27^, was verified in *M. schreibersii* with 100.0% editing efficiency. Functional overrepresentation analysis of the 2,125 recoded genes (Figure 5D; Supplementary Table 7) revealed significant enrichment in transmembrane receptor protein tyrosine kinase signalling, calcium ion transmembrane transport, and axon guidance. Key innate antiviral and inflammatory regulators undergo substantial recoding, including master type-I interferon factor IRF5 (up to 57.6% editing), restriction factors, janus kinase JAK2, antiviral GTPase MX1, and cytokines/receptors.

Genes with 3’ UTR editing (4,307 genes) showed marked enrichment in miRNA-mediated gene silencing, type II interferon-mediated signalling, and immune transcriptional activation. These results establish that A-to-I RNA editing in *M. schreibersii* operates as a dual-layer regulatory mechanism, expanding th structural repertoire of flight- and neural-related effectors while post-transcriptionally fine-tuning antiviral and inflammatory thresholds.

A fraction was used to train our BiLSTM neural network. After training our model with these events (50nt-flanking windows), we achieved an accuracy of 86.5% (Figure 6). If we compare it to the accuracies achieved with other species, it is near identical to the one obtained in mice using a specifi RNA-editing database for training (Table 6).

**Fig. 6:**
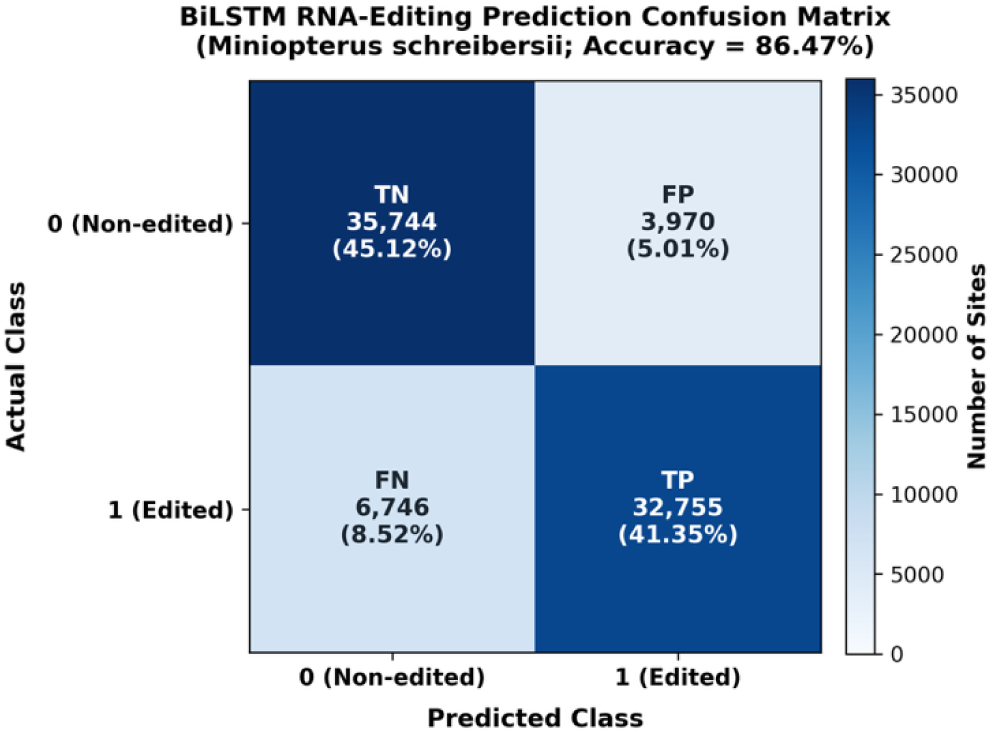
Accuracy of M. schreibersii RNA-editing prediction.

**Table 6.** RNA-editing prediction accuracies.

|  | *Miniopterus schreibersii* | *Canis lupus* | *Homo sapiens* | *Mus musculus* |
|---|---|---|---|---|
| Accuracies | 86% | 84% | 94% | 84% |

Using our cross-testing methodology^28^ (see methods) we can see how when cross testing 4 mammals (human, mouse, wolf and *Miniopterus*), only mouse and *Miniopterus* are able to consistently predict the other species with more than 60% accuracy. The other species, although achieving a similar or greater accuracy when predicting themselves, hover around 50% accuracy when predicting other species. This is strikingly curious in the case of the human trained algorithm as it achieves both the best and the worst accuracies obtained (Table 7).

**Table 7.** Cross-testing comparison.

| **Prediction**<br>**Training** | ***Miniopterus schreibersii*** | ***Canis lupus*** | ***Homo sapiens*** | ***Mus musculus*** |
|---|---|---|---|---|
| ***Miniopterus schreibersii*** | 86% | 68% | 63% | 62% |
| ***Canis lupus*** | 64% | 84% | 54% | 54% |
| ***Homo sapiens*** | 51% | 51% | 94% | 63% |
| ***Mus musculus*** | 63% | 68% | 74% | 84% |

## Discussion

Previous genomic investigations of *M. schreibersii* have historically been hindered by the limitations of short-read sequencing technologies. Over the last years only two assemblies of *M. schreibersii* have been published, one highly fragmented assembly with thousands of unassigned contigs. The other used long-read technology but still ended with ambiguous karyotypic assignments (misidentified as 2n = 44 linkage groups) ^21^. Here, by coupling native Oxford Nanopore ultra-long reads with high-throughput physical chromatin conformation capture, our assembly spans 1,767.14 Mb with a scaffold N50 of 83.06 Mb, assigning 98.5% of the genome into 23 distinct chromosome-scale scaffolds ( 2*n* = 46 ; autosomes SUPER_1 to SUPER_22, and chromosome X) alongside four resolved heterochromatic fragments of the Y chromosome (4.05 Mb) (Table 1, Figure 1).

In addition, the quality metrics achieved here fulfil the Earth BioGenome Project ^29^ (EBP) 7.7.Q53 benchmark, combining high base-level consensus accuracy ( QV = 53.29, *k* -mer completeness of 98.10%) with 96.7% complete single-copy BUSCO orthologs against vertebrata_odb10. Finally, our reconstructed mitochondrial genome (16,640 bp) provides a great maternal phylogenetic reference for future analysis. Compared not only to previous *Miniopterus* assemblies but to earlier Vespertilionoidea assemblies, this chromosome-scale reference genome establishes a new baseline for the family Miniopteridae, enabling precise synteny mapping, spatial chromatin architecture profiling, and accurate gene family orthology delineation, improving on the existing previous assemblies.

A striking topological feature revealed by our Hi-C contact map is the intense inter-chromosomal interaction network observed among the three smallest autosomes: SUPER_20 (27.86 Mb), SUPER_21 (17.96 Mb), and SUPER_22 (11.97 Mb). This trio displays prominent off-diagonal contact hubs connecting their termini to each other and to the pericentromeric regions of macrochromosomes (Figure 1). This raises some questions, as these micro-chromosomes could be technical Hi-C artifacts, dispensable supernumerary B-chromosomes, or an authentic evolutionary novelty of chiropteran nuclear architecture.

There are different ways to rule out technical or assembly artifacts. First, both the ONT ultra-long sequencing depth (116 x) and Omni-C physical contact distributions across SUPER_20–22 are uniform, exhibiting standard intra-chromosomal decay curves and uncompromised scaffold boundaries without chimeric misjoins. Second, unlike mammalian B-chromosomes, which are typically heterochromatic, gene-poor, repeat-dense, and genetically unstable, SUPER_20, SUPER_21, and SUPER_22 represent highly euchromatic A-chromosomes. They harbour the lowest repetitive element content across the entire genome (22.2%–23.5%), the highest GC content (46.53% to 49.67%, with SUPER_22 reaching the genome-wide apex), and more than double the transcript density of other macrochromosomes (61.5 to 87.8 transcripts/Mb vs. 42.4 transcripts/Mb on SUPER_1–9). They encode 4,523 functionally active loci, including indispensable cell cycle and immune effectors such as *CHEK1* on SUPER_20 and *IL16* on SUPER_21.

This means that, rather than an assembly artifact or B-chromosome signature, their elevated inter-chromosomal connectivity reflects two conserved biophysical properties of mammalian interphase 3D genome organization ^30,31^. In mammalian interphase nuclei, pericentromeric heterochromatin from acrocentric and telocentric chromosomes physically coalesces into punctate subnuclear structures termed chromocenters. In proximity ligation assays, this physical aggregation generates prominent, recurrent off-diagonal contact frequencies among non-homologous centromeric and telomeric compartments, as can be observed in *Miniopterus*. Furthermore, chromosome territories are non-randomly organized along a radial axis according to transcriptional activity and GC density: small, gene-dense, GC-rich chromosomes preferentially localize to the transcriptionally permissive nuclear interior, whereas large, gene-poor, AT-rich macrochromosomes are tethered toward the nuclear lamina at the periphery ^32,33^. The shared spatial occupancy of SUPER_20–22 within the central nuclear interior places them in constant physical proximity, creating the dense inter-chromosomal contact matrix captured by Hi-C.

Consequently, these micro-autosomes represent an authentic structural feature of the *M. schreibersii* karyotype, maintaining an intensely euchromatic and gene-dense chromatin environment segregated within the interior nuclear space.

Using CAFE5 to model phylogenomic birth-and-death events across 20 mammalian proteomes initially identified 151 candidate expansions and 565 contractions in *M. schreibersii*. However, when we analyse those results, we realised that automated gene family analysis in high-coverage transcriptomic assemblies frequently suffers from false-positive inflation caused by redundant transcript isoforms (e.g., StringTie models assigned distinct coordinates) and reference database asymmetries. Our spatial genomic curation found that 73 of the 151 expansion candidates were technical artifacts, (mainly split IGHV/IGLV and TRBV gene segments) reducing the dataset to 78 *bona fide* expanded families (Table 2, Figure 2).

Recent work by ^34^ revealed that vespertilionid bats possess two distinct IGH loci on separate chromosomes (A-IGH on chromosome 5 and B-IGH on chromosome 24 in *Eptesicus fuscus*), an inter-chromosomal duplication that originated after the divergence from Miniopteridae (∼37.5 Ma). However, in *M. schreibersii*, we find no evidence for an inter-chromosomal IGH duplication.

Among the 78 curated expansions, immune-related gene families reveal an evolutionary strategy that increases lymphocyte survival while tempering hyperinflammation. The former is represented with the expansion of the GTPases of immunity-associated proteins family (*GIMAP1*, *GIMAP2*, *GIMAP5B–D*; OG0000139, +4 copies). GIMAP proteins govern mitochondrial integrity and apoptotic thresholds in developing and mature T and B lymphocytes ^35,36^. Their expansion may be associated with long-term lymphocyte survival and immune memory maintenance in a species that encounters high viral diversity in highly dense colonies. In addition, there is a duplication of the tripartite motif (TRIM) E3 ubiquitin ligases (*TRIM60*, *TRIM61*, *TRIM75*; OG0000101, +3 copies), which recognize retroviral capsids and enveloped RNA virus replication complexes, targeting them for proteasomal degradation ^37^. This may be helping bats go through infections asymptomatically, a trait that bats are well-known for ^38^.

In striking symmetry, contracted gene families are highlighted by the reduction of NLRP inflammasome sensors (*NLRP1*, *NLRP5*, *NLRP14*; OG0000036, -6 copies). NLRP inflammasomes are primary drivers of caspase-1 activation, pyroptosis, and massive pro-inflammatory cytokine secretion (IL-1*β*, IL-18) ^14,39^. Their contraction in *M. schreibersii* aligns directly with the bat viral tolerance paradigm observed in other lineages ^40,41^, providing a mechanism that may dampen the risk of lethal cytokine storms during viral infection. Furthermore, contractions in the type I interferon locus (*IFNB1*, *IFNK*; OG0000024, -3 copies) reflect the evolutionary transition toward a constitutively expressed, low-level baseline antiviral state rather than an acute, hyper-inducible interferon surge ^42^.

Endogenous viral elements provide both a molecular fossil record of historical viral infections and a reservoir of raw genetic material for evolutionary novelties ^43,44^. Homology searches against viral databases (detectEVE) systematically flag conserved host exons whose viral homologs were historically acquired via host-to-virus horizontal gene transfer (HGT) ^22^. In *M. schreibersii*, domain and synteny auditing demonstrated that initial hits mapping to coding exons of interleukin-10 (EVE169), polyubiquitin (EVE023/024 and EVE040/041), and the DnaJ co-chaperone (EVE044) represent endogenous mammalian exons cross-matching viral virokines (poxviral/herpesviral vIL-10) ^45,46^, giant virus polyubiquitins ^47^, and cytopathic pestivirus *Jiv90* recombinants ^48,49^, alongside GC-rich CpG islands cross-matching proline-rich *Rubivirus* regions. Thus, rigorous filtering of these reciprocal mimics is essential to avoid misinterpreting viral insertions.

Nonetheless, focusing on the seven curated non-retroviral EVEs reveals a rich history of chiropteran interactions with single-stranded RNA viruses (Negarnaviricota: Bornaviridae and Mononegavirales), parvoviruses (Cossaviricota), and Preplasmiviricota ^50,51^. While intergenic integrations (EVE003, EVE098, EVE078, EVE079) are transcriptionally silenced across our available adult tissues, integrations residing within host gene models are actively transcribed and maintained across evolution. Regarding conservation, the intronic Mononegavirales L-polymerase element EVE029 within *LRIG2* and the Preplasmiviricota element EVE236 within the *BLOC1S6–SQOR* are shared syntenically with *M. myotis* (predating the divergence of Miniopteridae and Vespertilionidae 45 MYA). However, there is a key difference between these two EVEs. EVE029 has diverged at a neutral intronic rate (81.1% identity) ^52^ but EVE236 shows a 96.04% identity, highlighting the 303nt intronic region flanked by 81-83% identity areas, exposing a selective pressure to maintain the viral-like element. In addition, its association with the *BLOC1S6* (pallidin) locus, a core subunit of the BLOC-1 complex governing endolysosomal biogenesis and cytotoxic granule trafficking in natural killer (NK) and T cells, suggests potential cis-regulatory or non-coding functional exaptation in metabolic and lymphoid organs.

On the other hand, the exonic Bornaviridae P40 nucleoprotein integration EVE204 (BDV_P40) within the 3’ UTR of *TBL2* exhibits a clean empty pre-integration site in *M. myotis*, pinpointing a lineage-specific acquisition in *Miniopterus*. The co-transcription of EVE204 within the mature 3’ UTR of *TBL2* across 23 of 24 tissue libraries (reaching 17.80 TPM in testis and 14.90 TPM in lung) has direct functional implications for cellular stress and antiviral defense. *TBL2* encodes an endoplasmic reticulum (ER)-resident protein that binds phosphorylated PERK (*EIF2AK3*) under ER stress and unfolded protein response (UPR) conditions, regulating eIF2œ phosphorylation and stress-adaptive mRNA translation, a pathway central to both viral replication control and flight-induced proteotoxic stress ^53^. Furthermore, in mammals and bats, transcribed endogenous bornavirus-like nucleoprotein (*EBLN*) elements integrated into 3’ UTRs or introns have been shown to regulate host mRNA stability in *cis* or generate antisense piRNAs/siRNAs in *trans* that restrict exogenous bornavirus replication, particularly in germline (testis) and mucosal barrier (lung, intestine) tissues ^50,54^. Thus, the exonic integration of EVE204 into *TBL2* exemplifies how non-retroviral RNA virus sequences can be co-opted into the regulatory architecture of host stress-response genes.

While non-coding RNAs are nowadays an active field of research ^55,56^ most of the chiropteran comparative genomics have been focused on the protein-coding fraction of the genome ^14,57,58^. Thus, the non-coding portion of the transcriptome represents a regulatory layer, the understanding of which can better our comprehension of the immunological adaptations bats are well known for. This is especially true if we consider the extensive amount of examples where non-coding RNAs act as regulators of the immune system ^59–62^. Using our already tested filtering pipeline ^24,63^, we were able to focus on the long non-coding RNA portion and properly identify and categorise all the lncRNA present in our transcriptome (Figure 4).

Due to the lack (or rather low) selective pressure placed upon the nucleotidic sequence, lncRNAs are prone to undergo rapid primary sequence divergence ^64–66^. This makes conventional BLAST-based sequence alignment across species from different orders largely ineffective ^67,68^. However, as seen in Herrera-Ubeda et al.^24^, microsynteny can be used to unearth conserved long intergenic non-coding RNAs across great evolutionary distances. Using lincOfinder, we saw that across the ∼96 million years of mammalian divergence between *M. schreibersii* and *H. sapiens*, 4106 of the 5080 lincRNAs (80.83%) reside in conserved syntenic blocks and have an annotated human lncRNA ortholog (Supplementary Table 5). Compared to previous works, this is a strikingly high number, even for closely related species ^69^, but could be explained by the small number of tissues available for RNAseq, as well as the lack of any embryonic stages. In our transcriptome, only the lincRNAs with a basal expression on the 9 tissues analysed will be detected, and because of the low rate of boreoeutherian chromosomal rearrangements (Damas et al. 2022, Supplementary Figure 1), most of the lincRNAs will fall upon a conserved syntenic block.

Screening these syntenic blocks against curated mammalian immunome repositories revealed that 360 lincRNAs (8.77%) directly flank key immune effectors. Given that canonical immune genes comprise only 1.71% of all coding loci in *M. schreibersii* (mean syntenic block size = 3.89 genes), the null random expectation is 6.64% (272.8 expected syntenic blocks). The observed enrichment represents a statistically significant 1.32-fold increase over the genomic background. This overrepresentation could be linked to our aforementioned lincRNA discovery bias: Only the most basal expressed lincRNAs end up being detected, and as the bat immune system seems to be in a perpetual antiviral balance, its regulation via lincRNAs is constantly active, thus allowing us to sequence them. Of course, a lincRNA being in a conserved microsyntenic block with a gene does not directly mean that said lincRNA is regulating the genes on that block, but can be playing a role in the regulation of a related pathway ^71,72^.

A-to-I RNA editing catalyzed by ADAR deaminases provides a post-transcriptional mechanism to alter transcript sequences, coding potential, micro-RNA targeting affinity and secondary structures without altering the germline genome ^73–75^. Identifying the *M. schreibersii* editome across nine tissue libraries yielded 234,736 high-confidence unique editing events (≥ 10 x depth), 99.91% of which are canonical A-to-I transitions (Table 5, Figure 5). The overwhelming majority of editing sites reside in non-coding compartments: 65.28% intronic and 16.19% intergenic, localized within inverted repeat elements (SINEs and LINEs) in nascent transcripts (Figure 5A). In mammals, unedited double-stranded RNA (dsRNA) hairpins formed by inverted repeats are recognized by cytosolic pattern recognition receptors (MDA5, RIG-I, PKR), triggering aberrant type-I interferon production and autoimmune pathology ^76^. Pervasive adenosine deamination destabilizes these dsRNA duplexes via A-to-I mismatches, preventing auto-inflammatory sensing. In *M. schreibersii*, this dense non-coding editome likely acts as an intrinsic threshold buffer, preventing self-nucleic acid sensing.

When switching our focus to the editing within open reading frames, we identified 3,715 nonsynonymous recoding events (1.58% of the editome) across 2125 unique genes. One of them was the canonical neuroprotective recoding site, the Q/R site (Q607R) within the pore of the AMPA glutamate receptor *GRIA2* (SUPER_4:47,661,840), which was verified at 100.0% editing efficiency ^25,27^.

Functional enrichment analysis showed that recoded transcripts are significantly enriched in Transmembrane Receptor Protein Tyrosine Kinase Signaling, Calcium Ion Transport, and Axon Guidance Pathways among others. But if we investigate the events targeting immune related genes, key innate antiviral regulators undergo substantial nonsynonymous recoding. These include the master type-I interferon transcription factor IRF5 (K31E up to 57.6% editing), tripartite motif restriction factor TRIM62 (S189G at 54.8%), janus kinase JAK2 (D370E and K857E), and the antiviral GTPase MX1 (N641D). Furthermore, 3’ UTR editing heavily targets miRNA machinery (*AGO2*, *AGO3*, *AGO4*, *TNRC6A/B*) and type II interferon signaling components (*IFNGR1/2*, *IRF1*, *TYK2*). All of these extensive recoding points to a scenario where, rather than relying solely on fixed genetic mutations, RNA editing provides *M. schreibersii* with dynamic epitranscriptomic plasticity, expanding the structural repertoire of metabolic and neural effectors while fine-tuning the activation thresholds of antiviral signaling cascades.

Using our bidirectional LSTM (BiLSTM) network with attention layer trained on 50+50nt sequence and structural contexts, the model achieved an internal predictive accuracy of 86% in *M. schreibersii* (Figure 6, Table 6).

In cross-species evaluation across four mammals (*Miniopterus*, mouse, wolf, and human), an intriguing evolutionary pattern emerged (Table 7): Models trained on *M. schreibersii* and *Mus musculus* reciprocally cross-predicted editing sites in each other with 62%–68% accuracy, consistently outperforming cross-predictions involving human or wolf algorithms. Conversely, while human-trained models achieved high intra-species accuracy (94%), they exhibited poor cross-predictive generalization to non-primate mammals (51% in bat and wolf), and wolf-trained models hovered near random chance (54%) when predicting mouse or human.

This reciprocal cross-predictive fidelity between *Miniopterus* and mouse reveals that despite >90 million years of divergence, bats and rodents share a deeply conserved local cis-regulatory grammar and structural characteristics that direct ADAR deamination. The deamination machinery in bats does not rely on novel sequence rules but rather leverages an ancestral mammalian structural code with elevated efficiency to orchestrate transcriptomic plasticity.

## Conclusions

By integrating a contiguous chromosome-scale genome assembly with multi-tissue transcriptomics, long non-coding RNA synteny, endogenous viral element mapping, and same-individual based RNA editomics, this study provides a comprehensive multi-omic framework for the common bent-wing bat (*Miniopterus schreibersii*).

Our findings not only provide an improved version of the *Miniopterus* genome, but they also demonstrate that chiropteran viral tolerance does not stem from a single genomic innovation, but rather from the coordinated convergence of multiple evolutionary layers. First, our curated phylogenomic analysis revealed 78 bona fide expansions balancing intense flight metabolism and ferroptosis defence against immune restraint. In addition, unlike vespertilionids, *Miniopterus* immune diversification does not rely on IGHV/IGLV chromosomic duplications, but rather on targeted expansions in lymphocyte survival GTPases alongside adaptive contractions in pro-inflammatory *NLRP* inflammasomes and type I interferons.

Then, the search of paleoviral insertions validated seven nrEVEs, highlighted by a *Miniopterus*-specific Bornaviridae nucleoprotein integrated directly into the 3’ UTR exon of the ER-stress regulator *TBL2* and actively expressed across several tissues, with another integration of a Preplasmiviricota fragment showing signs of purifying selection in *Myotis* and *Miniopterus*.

When looking at the non-coding fraction, we identified deep microsyntenic conservation of 4514 lincRNAs across ∼96 million years of divergence, significantly enriched at core immune loci, pointing to conserved topological chromatin scaffolds that have a role regulating the immune transcription in the *Miniopterus* permanent antiviral state. This is coupled with the massive editome dominated by non-coding A-to-I deaminations shielding host cells from sterile dsRNA auto-inflammation, in addition to nonsynonymous recoding in core antiviral factors and deep cross-species conservation of editing sequence rules between bats and rodents.

Together, these results establish *M. schreibersii* as a definitive Palearctic reference model for mammalian viral tolerance, metabolic adaptation, and host-pathogen coevolution. As specialized flyers, long-lived mammals, and asymptomatic viral hosts, bats remain truly battastic creatures.

## Methods

### Specimen Sampling, High-Molecular-Weight DNA Extraction, and Sequencing

A wild male specimen of the *M. schreibersii* (sample ID BA1092 / 9911AH, ToLID mMinSch1) was collected from a spring colony in Garraf Park (Catalonia, Spain), adhering strictly to regional and institutional ethical frameworks for wildlife research. High-molecular-weight (HMW) genomic DNA was extracted from flash-frozen tissue and prepared for long-read sequencing using the Oxford Nanopore Technologies (ONT) Ligation Sequencing Kit (SQK-LSK114). Ultra-long native DNA sequencing was performed on an ONT PromethION platform across two R10.4.1 flow cells (PAO37309 and PAQ19246) at the Centro Nacional de Análisis Genómico (CNAG, Barcelona, Spain). Basecalling was executed using Guppy v6.5.7 with the super-accuracy model (dna_r10.4.1_e8.2_400bps_5khz_sup.cfg), generating 205.14 Gb of long-read data (12.07 million reads; read N50 = 34.33 kb; maximum single-read length = 1.25 Mb), representing approximately 116 x nuclear genome coverage.

In parallel, proximity ligation libraries were constructed using the Dovetail Omni-C protocol (sample ID BA5956 / “bat cat 2”) to capture long-range physical chromatin contacts and short-read sequence data. Omni-C libraries were sequenced on an Illumina platform at CNAG, generating 550.04 million paired-end reads (166.11 Gb yield), providing approximately 94 x physical coverage for chromosome conformation scaffolding and short-read base polishing.

### Genome Assembly, Polishing, Chromosome-Scale Scaffolding, and Quality Control

The nuclear genome was assembled following the European Reference Genome Atlas (ERGA) standard pipeline. Quality filtering was performed on ONT reads with Filtlong v0.2.1 and on Illumina paired-end reads with Trim_Galore v0.6.7 ^77^ (--gzip -q 20 --paired --retain_unpaired). *De novo* long-read contig assembly was conducted using NextDenovo v2.5.0 ^78^. Assembly polishing was performed with short reads using NextPolish v1.4.1 ^79^. Redundant haplotypic duplications were identified and purged using purge_dups v1.2.6 ^80^. Chromosome-scale scaffolding was performed with YaHS v1.2a ^81^ utilizing Dovetail Omni-C physical contact pairs.

The resulting scaffolded assembly underwent comprehensive manual curation and quality control using PretextView v0.2.5 ^82^ (14 interventions/Gb). Potential contamination was evaluated using BlobTools against NCBI taxonomic databases.

### Mitochondrial Genome Assembly and Annotation

The complete circular mitogenome of *M. schreibersii* was assembled from whole-genome ONT reads using the FOAM pipeline v0.5 (CNAG-AAT; https://github.com/cnag-aat/FOAM). In brief, ONT Q20+ reads shorter than 18,000 bp were bait-mapped against the *Miniopterus fuliginosus* reference mitogenome (NC_044489.1) using minimap2 v2.24 ^83^. Bait-selected reads were assembled *de novo* with Flye v2.9 ^84^ under metagenomic mode (--meta --scaffold -i 1), generating a circular contig of 16,961 bp. The contig was annotated with MITOS v2.1.3 ^85^ to identify the tRNA-Phe (*trnF*) locus and oriented to set its canonical origin at the start of *trnF* using orient_mitogenome_v1.pl. Following orientation, the mitogenome was polished with an iterative round of Flye using mitochondrial ONT reads to eliminate residual k-mer errors. The finalized 16,640 bp circular mitogenome was annotated with MITOS v2.1.3.

### Multi-Tissue RNA-Seq and Transcriptome Assembly

Total RNA was isolated from nine anatomical tissues (patagium/wing epithelium, heart, liver, intestines, spleen, brain, lungs, testis and skeletal muscle) across 24 paired-end libraries (multi-tissue biological replicates) from three sequenced Catalonian specimens. RNA-seq libraries were sequenced on an Illumina NovaSeq 6000 platform ( 2 x 150 bp). Quality-filtered reads were aligned against the soft-masked chromosome assembly using STAR v2.7.10a ^86^ in two-pass multi-sample mode (--outSAMstrandField intronMotif), pooling splice junctions discovered across all 24 libraries during Pass 1 prior to second-pass alignment. Transcript models were assembled per library and merged into a non-redundant reference transcriptome using StringTie v2.2.1 ^87^. Protein-coding regions were predicted using TransDecoder v5.5.0 ^88^ guided by Pfam and UniProt database searches.

### Gene Family Dynamics and Evolutionary Modeling (CAFE5)

Comparative genomic analysis was performed across 20 representative vertebrate proteomes: one outgroup (*Gallus gallus*), one metatherian (*Monodelphis domestica*), one afrotherian (*Loxodonta africana*), two euarchontoglirans (*Homo sapiens*, *Mus musculus*), six non-bat laurasiatherians (*Sorex araneus*, *Canis lupus familiaris*, *Felis catus*, *Equus caballus*, *Bos taurus*, *Sus scrofa*), three yinpterochiropterans (*Rhinolophus ferrumequinum*, *Rousettus aegyptiacus*, *Hipposideros armiger*), and six yangochiropterans (*Miniopterus schreibersii*, *Myotis myotis*, *Pipistrellus kuhlii*, *Eptesicus fuscus*, *Desmodus rotundus*, and *Molossus molossus*).

Canonical protein isoforms (longest ORF per locus) were clustered into orthologous groups (orthogroups) using OrthoFinder v3.1.2 ^89^ with Diamond v2.1.9 under ultra-sensitive mode (-S diamond_ultra_sens) and MSA-based species tree reconstruction (-M msa). An ultrametric chronogram was constructed in R using penalized likelihood in ape::chronos v5.8, calibrated with nine secondary fossil constraints from TimeTree (timetree.org): Amniota root (312–326 Ma), Mammalia root (155–165 Ma), Theria root (150–160 Ma), Placentalia (95–105 Ma), Boreoeutheria (92–99 Ma), Laurasiatheria (78–86 Ma), Chiroptera crown (58–66 Ma), Yinpterochiroptera crown (50–58 Ma), and Yangochiroptera crown (45–52 Ma).

Gene count matrices were filtered to exclude families with extreme lineage-specific expansions ( ≥ 100 genes in any single species) or restricted to a single taxon. Gene family birth-and-death rates were estimated using CAFE v5.1.0 ^90^. Genome assembly and annotation errors were modelled using the -e flag, estimating an error parameter *ε* = 0.0129. Global birth-death rates (*λ* = 0.000296 substitutions/Myr) were estimated under a homogeneous birth-death model conditioned on c. Branch-specific p-values were calculated via Monte Carlo simulations (1,000 iterations), designating families with *p* < 0.05 on the *M. schreibersii* terminal branch as significantly expanded or contracted. Candidate expansions were subjected to rigorous case-by-case spatial genomic curation to eliminate overlapping transcript models and reference database asymmetries, establishing a definitive set of *bona fide* expanded families. Functional enrichment of curated expansions and contracted families was assessed using Enrichr ^91^ (Fisher exact test against the library gene universe, Benjamini-Hochberg FDR correction; only named genes were queried) across Gene Ontology Biological Process (GO BP 2023), KEGG Pathways (2021 Human), and Reactome (2022).

### Endogenous Viral Element (EVE) Mining and Tissue Expression Quantification

Endogenous viral elements were identified using detectEVE ^92^ against the soft-masked chromosome assembly. To eliminate false positives arising from ancient host-to-virus horizontal gene transfer (viral mimicry of cellular genes) or compositional bias in GC-rich simple repeats, non-retroviral EVE (nrEVE) candidates were translated in all six reading frames and scanned against Pfam-A v35.0 using hmmscan (HMMER v3.3.2), cross-referenced against vertebrate RefSeq proteomes (BLASTX), and aligned with 10-kb flanking genomic windows against orthologous loci in *Myotis myotis* (HLmyoMyo6-2.fa) using BLASTN to verify empty pre-integration sites or shared ancestral synteny.

### lncRNA Discovery Pipeline and Positional Microsynteny Analysis

Long non-coding RNAs were identified through an integrative four-step computational pipeline:

1. **Length filtering:** All reconstructed transcript isoforms were filtered to retain models ≥ 300 nt.
2. **Coding potential filtering:** Candidate transcripts were evaluated with CPAT v3.0.4 (Wang et al. 2013), discarding models with a coding probability ≥ 0.4.
3. **Sequence homology filtering:** Transcripts were searched against the NCBI non-redundant protein database (nr) using BLASTX (E-value cutoff 1 x 10^−4^) and against the Pfam-A v35.0 database using HMMER v3.3.2 (E-value cutoff 1 x 10^−4^); any transcript showing significant coding domain homology was excluded.
4. **ORF length filtering:** Potential ORFs were predicted with TransDecoder; models containing ORFs ≥ 150 amino acids were eliminated. The surviving putative lncRNAs were classified into intergenic, antisense, intragenic, and bidirectional based on genomic orientation relative to protein-coding loci.

Positional microsynteny between *M. schreibersii* and *Homo sapiens* (GRCh38) was established using LincOFinder ^24^. Syntenic blocks were defined by conserved gene order and orientation of upstream and downstream coding gene anchors flanking the focal lncRNA (intergenic and bidirectional). Conserved blocks were cross-referenced against NCBI human RefSeq annotations to identify homologous human non-coding genes. Syntenic blocks were screened against curated mammalian immune genes (ImmPort and GO:0002376 “immune system process”). Statistical enrichment of immune-associated syntenic blocks over the genomic background (1.71% immune coding genes; 3.89 genes/block) was tested using exact Poisson-binomial tests.

### RNA-Editing Identification, Functional Annotation, and BiLSTM Deep Learning

To profile the *M. schreibersii* RNA editome, nine stranded paired-end RNA-seq libraries (specimen 3) prepared from diverse tissues of the sequenced individual were aligned against the chromosome assembly using Bowtie2 v2.4.5 ^94^ in local alignment mode. Matched whole-genome DNA sequencing reads from the same Catalonian individual were aligned with Magic-BLAST v1.5.0 ^95^ to generate an empirical genomic DNA baseline. Alignments were processed, sorted, and deduplicated using SAMtools v1.17 ^96^.

Variant calling was executed across RNA alignments using BCFtools mpileup and call (v1.17) ^96^ with base quality recalibration (min-BQ 20, min-MQ 30). Candidate RNA variants (A-to-G or T-to-C variants that could represent A-to-I editing) were filtered against the matched DNA baseline, retaining sites only if: (i) the locus possessed zero evidence of variant alleles in the DNA baseline, (ii) total RNA sequencing depth was ≥ 10 x in at least one tissue library, and (iii) the editing frequency was ≥ 2% in at least one library.

Functional consequences were determined by intersecting editing coordinates with the assembled transcriptome. For sites falling within open reading frames (CDS), phase and codon positions were extracted, and reference triplets were fetched from the soft-masked genome assembly using pysam. Reference and in silico edited codons (incorporating an inosine-directed G substitution on the mRNA strand) were translated using the standard genetic code to classify events as recoding (missense), synonymous (silent), or stop-altering. Sites outside CDS were partitioned into 5’ UTRs, 3’ UTRs, non-coding exons, introns, or intergenic intervals. Functional pathway overrepresentation was evaluated via Enrichr.

For neural network modeling, candidate sites and non-edited adenosines were centered within symmetrical 50-nt genomic windows (101 nt total) and one-hot encoded into a dual-channel matrix capturing sequence composition and secondary structural folding propensities. A bidirectional Long Short-Term Memory (BiLSTM) network with an attention layer was trained using 10-fold cross-validation. For cross-species generalization, models independently trained on *M. schreibersii*, *H. sapiens*, *M. musculus*, and *C. lupus* were cross-evaluated across all pairwise combinations without retraining ^28^.

## Data Availability

All the data generated from sequencing is available under the bioprject id PRJNA1559021. The fully assembled and annotated genome is available in ncbi as SAMN64834586 : mMinSch1 and all the transcriptomic and genomic raw data is uploaded to SRA.

## Acknowledgements (optional)

We would like to acknowledge Claudia Fernandez and all of Jordi Garcia’s lab members for the scientific discussion. M.Z.-A. is a recipient of an FPU (Formación del Profesorado Universitario) scholarship from the Spanish Ministerio de Universidades. J.G-F. received funding from the Ministerio de Educación y Ciencia (grant number PID2020-117820GB-I00 and PID2023-146794NB-I00).

