## Supplementary Figure 1 for "Elucidating Bat SuperImmunity: Insights from the Bent-Wing Bat Genome Assembly and Multi-Omics Analysis"

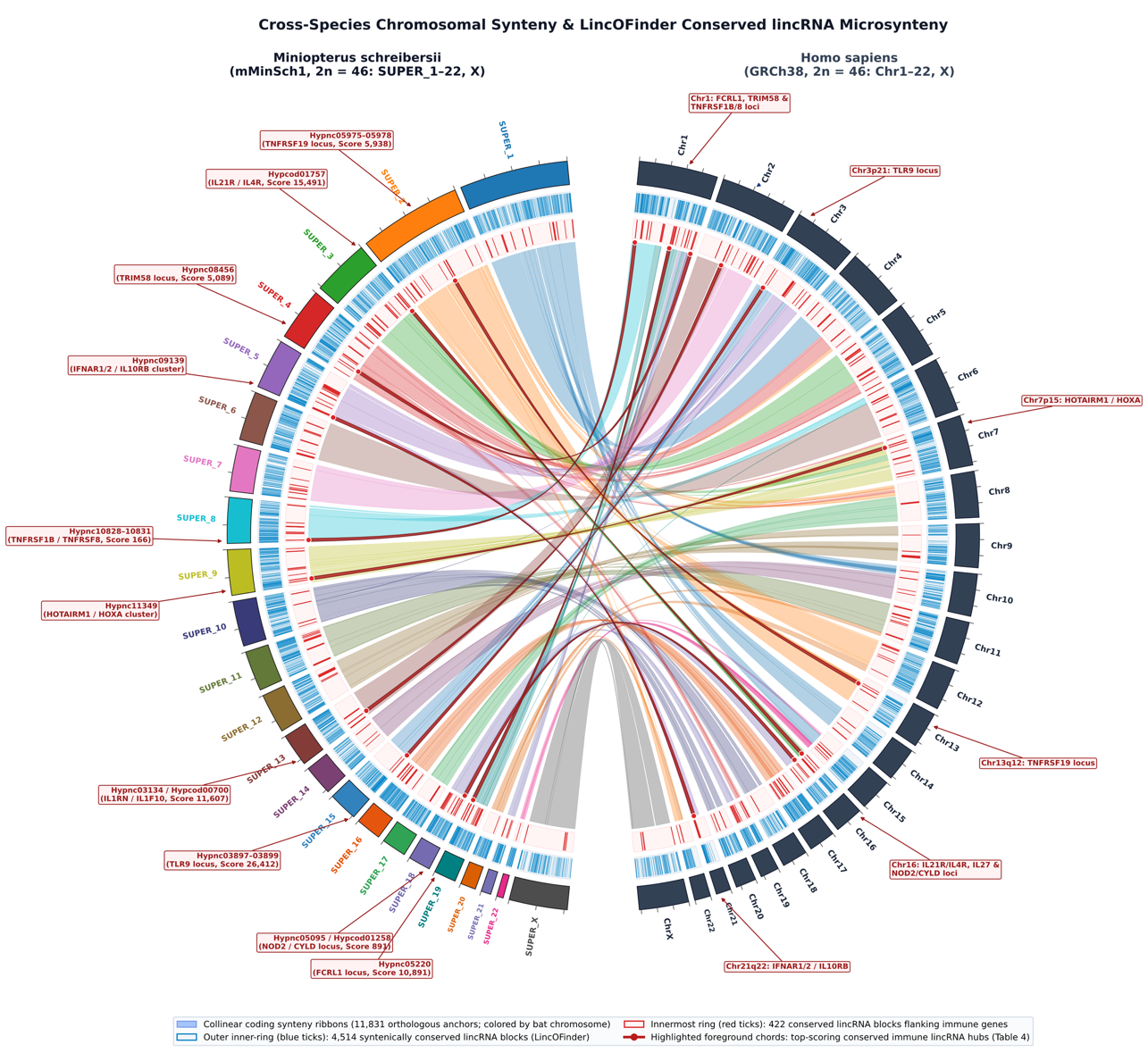
**Supplementary Figure 1: Cross-species chromosomal macrosynteny and LincOFinder conserved lincRNA microsynteny between Miniopterus schreibersii and Homo sapiens.**

Ideograms on the left semicircle represent the 22 autosomes (SUPER_1–SUPER_22) and X chromosome (SUPER_X) of M. schreibersii (mMinSch1, $2n=46$), while ideograms on the right semicircle represent human chromosomes (Chr1–Chr22 and ChrX, GRCh38, $2n=46$), with outer tick marks every 50 Mb. The outer concentric inner-track (blue ticks) displays the genomic distribution of the 4514 syntenically conserved lincRNA blocks identified by LincOFinder. The innermost concentric track (red ticks) highlights the conserved lincRNA blocks flanking immune-related genes (360 of which harbor an annotated human lncRNA ortholog). Interior translucent ribbons connect 132 collinear macrosyntenic blocks (11,831 orthologous protein-coding gene anchors), colored according to the M. schreibersii chromosome of origin. Bold crimson foreground chords and outer callout boxes highlight the top-scoring conserved immune lincRNA hubs summarized in Table 4.
